## Supplementary material for "3’HS1 CTCF binding site in human β-globin locus regulates fetal hemoglobin expression": Supplemt Figure 1-8

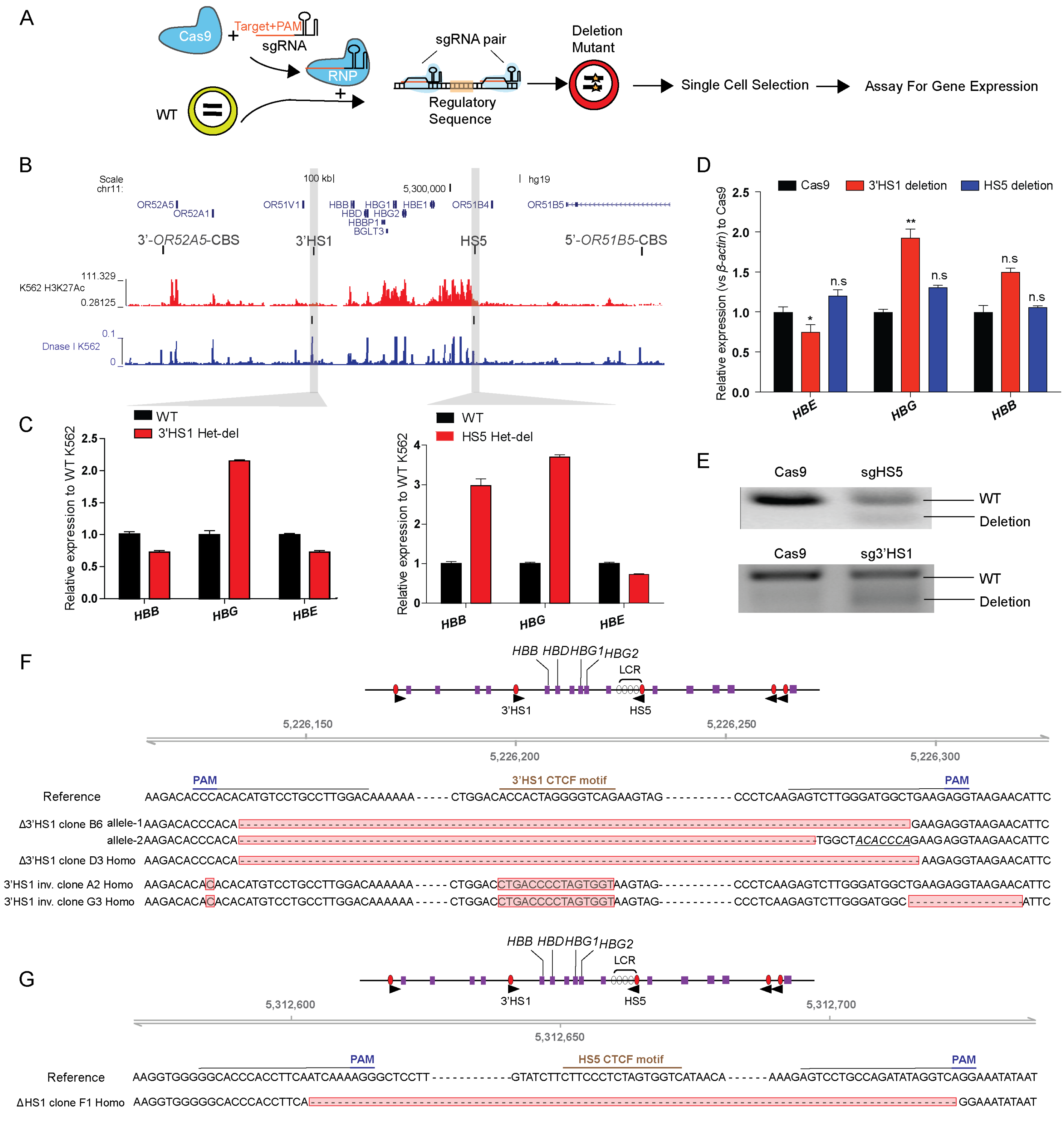


**Figure S1. CTCF binding site around β-globin gene cluster regulated β-globin gene expression.** (A). The experimental scheme of CTCF binding site deletion by CRISPR/Cas9. (B)**.** CTCF binding site and chromatin landscape (H3K27ac and Dnase hypersensitivity footprint is shown) around β-globin genes (C). The β-globin genes expression in the K562 clones with CBS deletion. (D). The β-globin genes expression in the bulk HUDEP-2 cells with HS5 and 3'HS1 deletions. N=3. mean±S.D is displayed. n.s not significant. * p<0.05, ** p<0.01. Two tailed t-test is performed. (E). Deletion fraction of 3'HS1 and HS5 in the bulk HUDEP-2 population tested in (D). (F-G)**.** The sanger sequencing validation result of Δ3'HS-1 clones and 3'HS-1 inversion clones as well as HS5 deletion clone (G). Homo indicates the deletion locus is homozygous. Insertion site is marked italic with underline text.

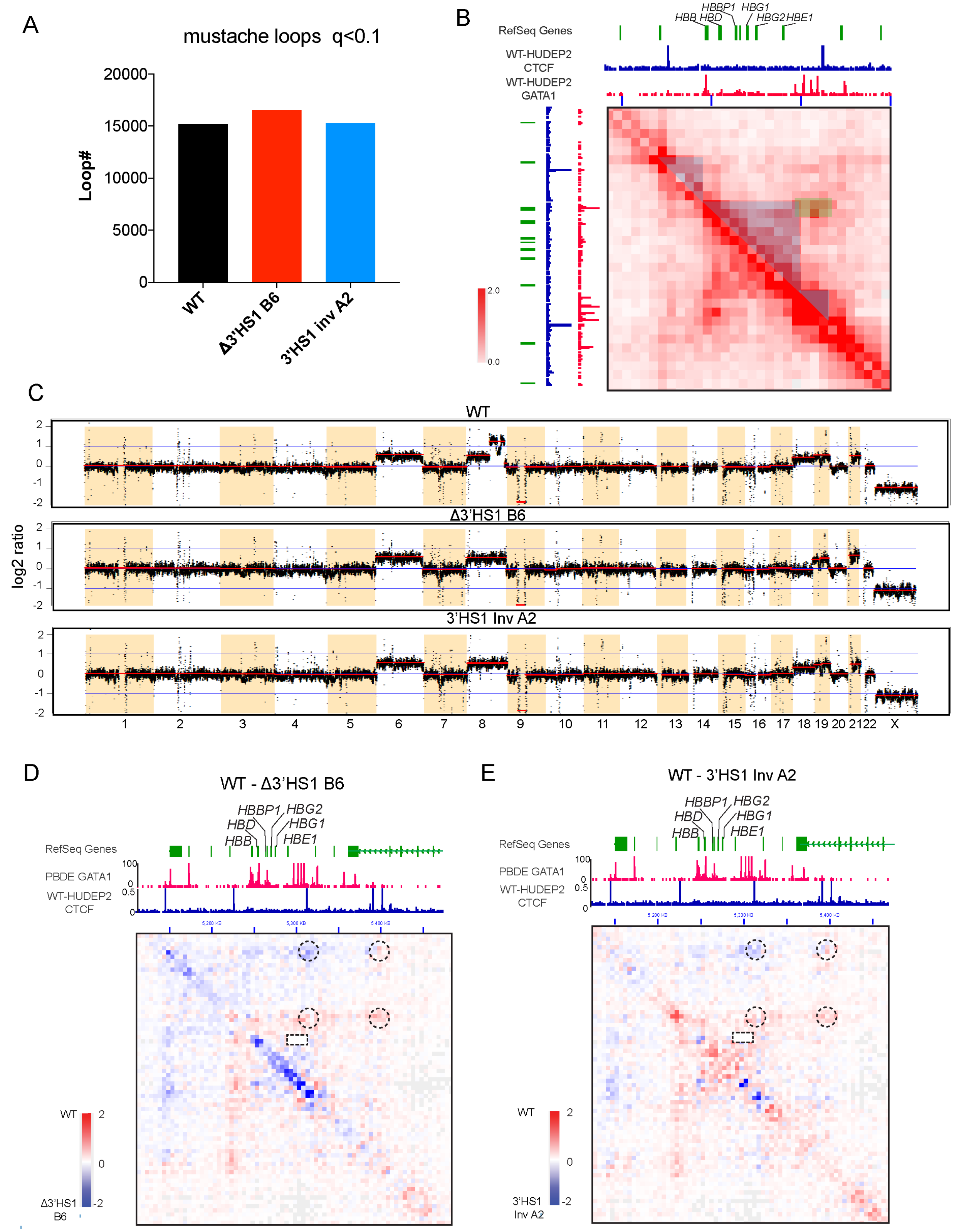


**Figure S2. 3D genomics change in Δ3'HS-1 clones and 3'HS-1 inversion HUDEP-2 cell clones.** (A). The loop number called by mustache with parameter of q < 0.1{ref}. (B). The 3D genomics interaction landscape in β-globin gene cluster. Blue shaded region indicates the sub-TAD domain between HS5 and 3'HS-1 CBS. Green shaded region covers the interactions between LCR (HS1-4) and *HBB* interactions. (C). The CNV profile of 3 cell clones inferred from HiC data by HINT{ref}. (D-E). The juicebox view of WT- Δ3'HS-1 clone B6 (D) and WT-3'HS-1 inversion clone A2 (E) contact map. The CBS associated chromosomal loops are circled out. LCR-HBB interaction is highlighted by rectangular box.

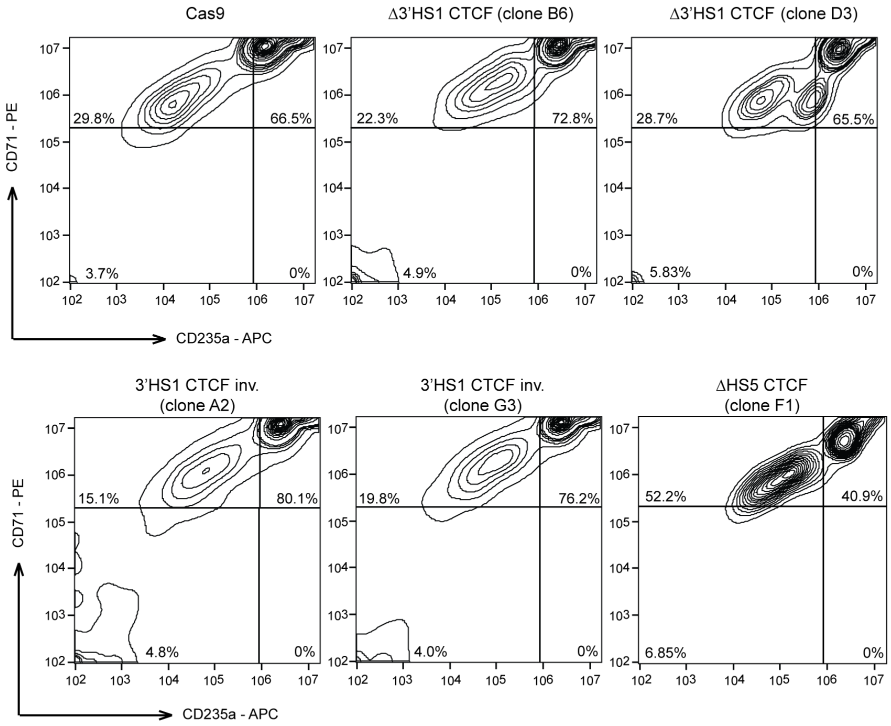


**Figure S3.** The differentiation stage of HUDEP-2 cell clones used in **Figure1** profiled by Flow Cytometry of CD71 and CD235a.

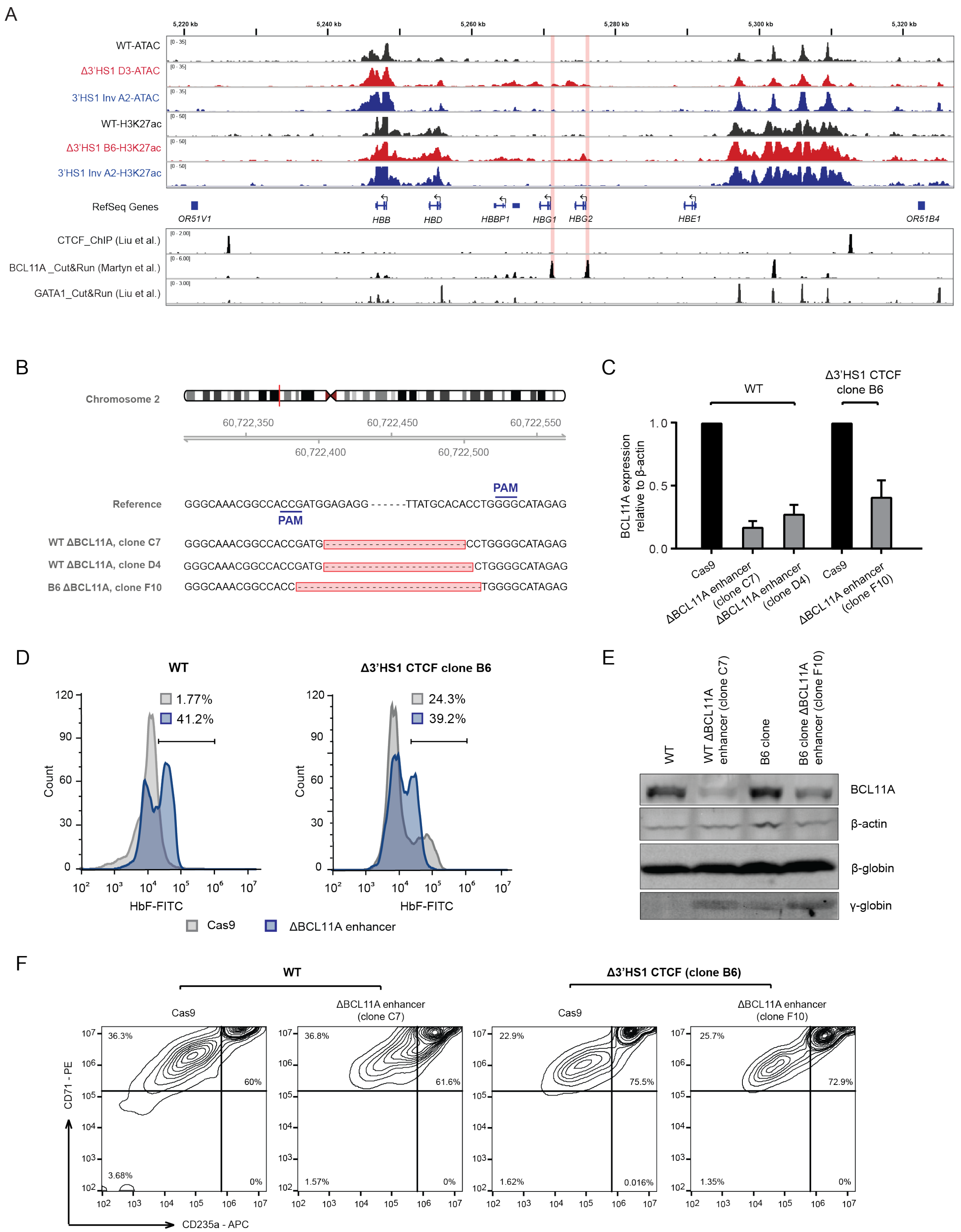


**Figure S4. BCL11A loss further promote fetal hemoglobin induction in Δ3'HS-1 background. (A)**. IGV track view of ATAC-seq and H3K27ac ChIP-seq at β-globin gene locus of Δ3'HS1 HDUEP-2 cell clones, 3'HS1 inversion HDUEP-2 cells and wild-type HUDEP-2 cells. GATA1, CTCF and BCL11A Cut&Run data is shown below the track. Regions highlighted in orange are paralogous *HBG1/2* promoter. (B). Sanger sequencing of *BCL11A* +58 enhancer disrupted clones. (C). qPCR quantification of *BCL11A* gene expression in +58 enhancer deleted clones. (D). Flow Cytometry measurement of HbF in WT and B6 clone after the deletion of *BCL11A* +58 enhancer by CRISPR/Cas9. (E). Western Blot quantification of BCL11A protein in *BCL11A* +58 enhancer deleted clones in WT and Δ3'HS-1 background. (F). The differentiation stage of *BCL11A* +58 enhancer disrupted HUDEP-2 cell clones profiled by Flow Cytometry of CD71 and CD235a.

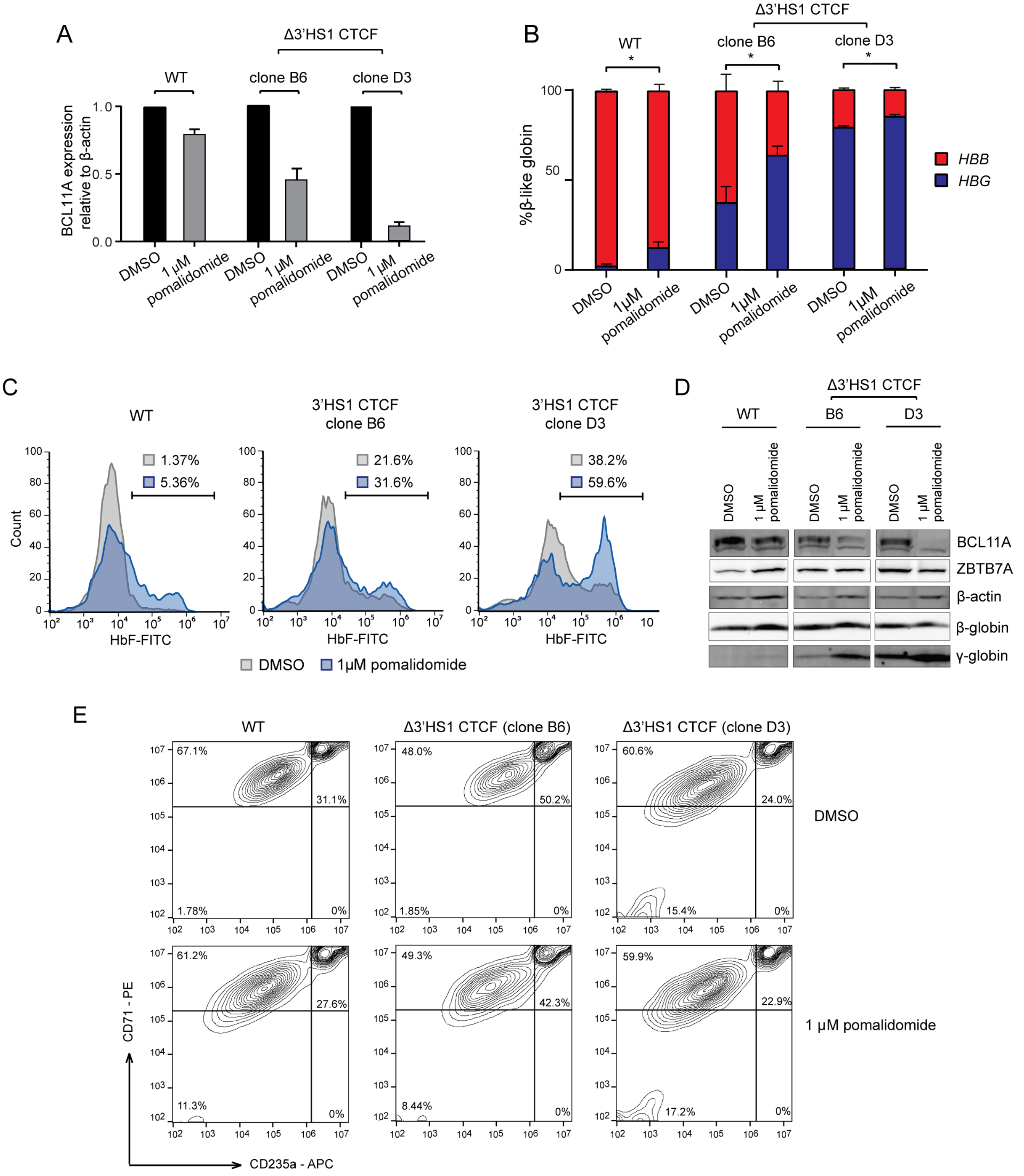


**Figure S5. Pomalidomide enhance fetal hemoglobin production induced by 3'HS-1 deletion** (A). qRT-PCR quantification of *BCL11A* in the Pomalidomide treated WT and 3'-HS-1 deleted HUDEP2 cell clones. (B). Composition of beta like globin by qRT-PCR in clones treated with DMSO and 1uM Pomalidomide described in panel (A). (C). The flow cytometry plot of HbF in clones treated with DMSO and 1uM Pomalidomide described in panel (A) (D). Western blot of BCL11A, ZBTB7A, β-globin and γ-globin clones treated with DMSO and 1uM Pomalidomide described in panel (A). (E). The differentiation stage of HUDEP-2 cell clones used in panel (A) profiled by Flow Cytometry of CD71 and CD235a.

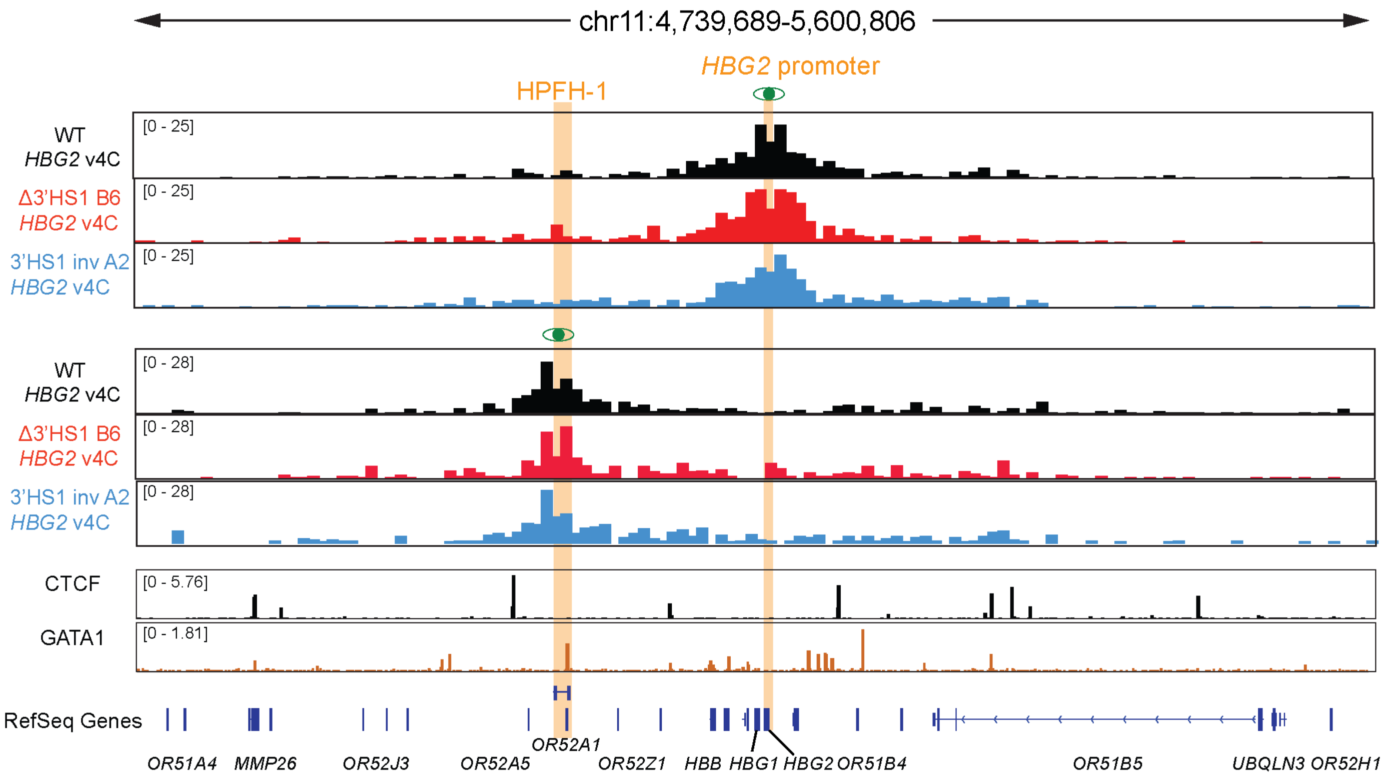


**Figure S6. *HPFH-1* and *HBG2* interactions in edited HUDEP-2 cells.** The v4C tracks generated by Juicebox from WT, Δ3'HS1 HUDEP2 clone B6 and 3'HS1 inversion clone A2. The *HPFH-1* region is highlighted in orange and viewpoint of 4C is highlighted in orange and eye symbol on *HPFH-1* enhancer and *HBG2* promoter region respectively.

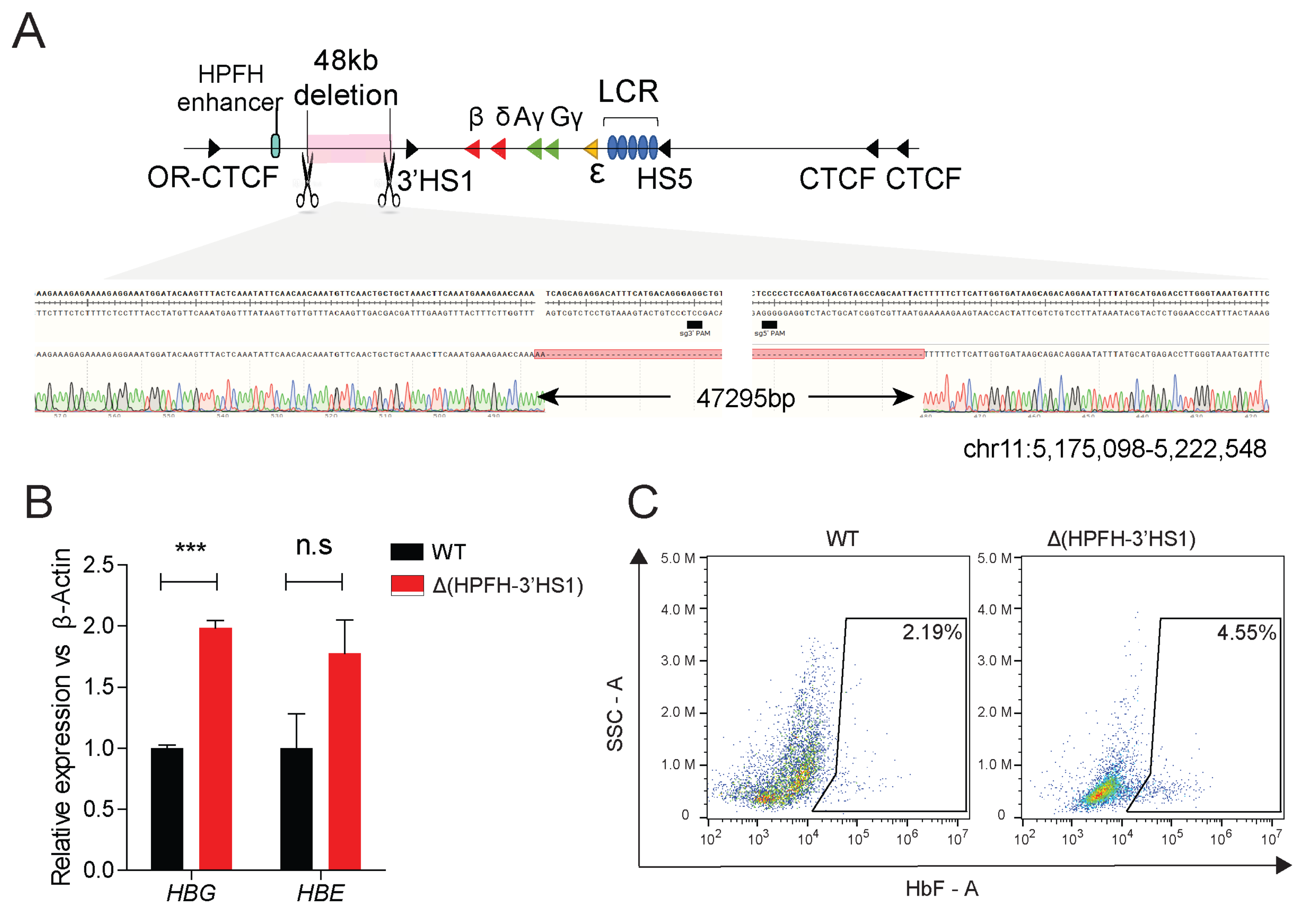


**Figure S7. Deletion between HPFH enhancer and 3'HS1 increase the *HBG1/2* and *HBE* expression** (A). The experimental scheme of deleting 48kb region between HPFH enhancer and 3'HS1. The sanger sequencing validation result of a Δ(HPFH-3'HS1) clone B5 is displayed in the lower panel. (B). *HBG* and *HBE* expression in WT and Δ(HPFH-3'HS1) clone B5. Mean ± S.D. is shown. ***: $p < 0.001$, n.s: not significant. t-test was performed to determine the p value. (C). The flow cytometry plot of HbF in WT and Δ(HPFH-3'HS1) clone B5.

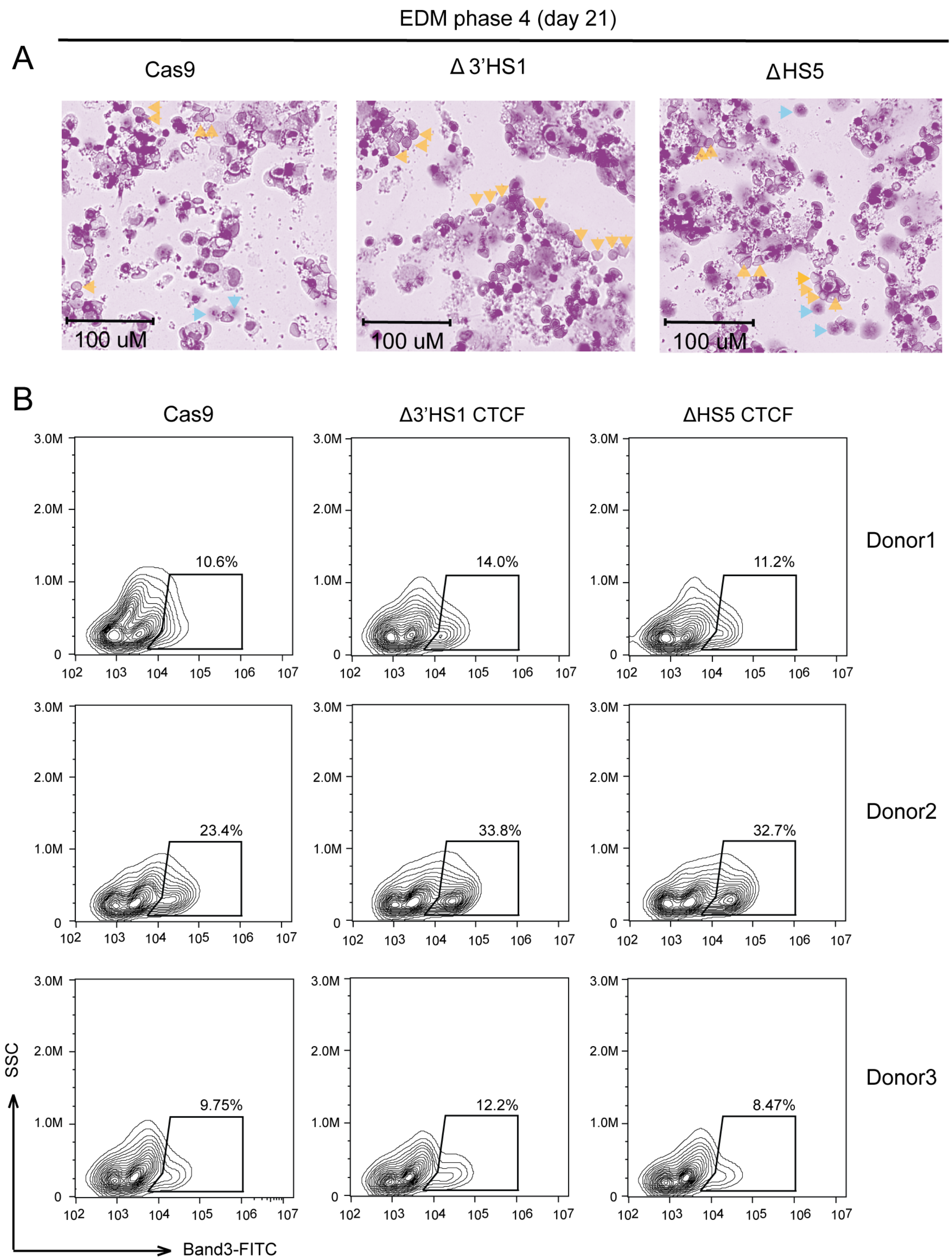


**Figure S8. 3'HS1 deletion in HSPC.** (A). Giemsa-Wright staining of differentiated erythroid cells from HSPC electroporated with Cas9, 3'HS-1 guide RNA pair and HS5 deletion guide RNA pair at differentiation culture day 21. Orange arrow indicates the enucleated red blood cells and blue arrows indicates the reticulocytes. (B). Differentiation stage of 3 HSPCs by flow cytometry of Band3 from different donors electroporated with Cas9, 3'HS-1 guide RNA pair and HS5 deletion guide RNA pair at differentiation culture day 21.
